# Compositional restrictions in the flanking regions give potential specificity and strength boost to binding in short linear motifs

**DOI:** 10.1101/2024.05.13.593809

**Authors:** Veronika Acs, Andras Hatos, Agnes Tantos, Lajos Kalmar

## Abstract

Short linear motif (SLiM)-mediated protein–protein interactions play important roles in several biological processes where transient binding is needed. They usually reside in intrinsically disordered regions (IDRs), which makes them accessible for interaction. Although information about the possible necessity of the flanking regions surrounding the motifs is increasingly available, it is still unclear if there are any generic amino acid attributes that need to be functionally preserved in these segments. Here, we describe the currently known ligand-binding SLiMs and their flanking regions with biologically relevant residue features and analyse them based on their simplified characteristics. Our bioinformatics analysis reveals several important properties in the widely diverse motif environment that presumably need to be preserved for proper motif function, but remained hidden so far. Our results will facilitate the understanding of the evolution of SLiMs, while also hold potential for expanding and increasing the precision of current motif prediction methods.

**Author summary:** Protein–protein interactions between short linear motifs and their binding domains play key roles in several molecular processes. Mutations in these binding sites have been linked to severe diseases, therefore, the interest in the motif research field has been dramatically increasing. Based on the accumulated knowledge, it became evident that not only the short motif sequences themselves, but their surrounding flanking regions also play crucial roles in motif structure and function. Since most of the motifs tend to be located within highly variable disordered protein regions, searching for functionally important physico-chemical properties in their proximity could facilitate novel discoveries in this field. Here we show that the investigation of the motif flanking regions based on different amino acid attributes can provide further information on motif function. Based on our bioinformatics approach we have found so far hidden features that are generally present within certain motif categories, thus could be used as additional information in motif searching methods as well.

## Introduction

Short linear motifs (SLiMs) are short modules of protein sequences that play crucial roles in mediating and regulating protein–protein interactions, where transient and dynamic binding is needed (e.g., signal transduction and regulatory processes) [1]. SLiMs are represented by a limited number of constrained affinity- and specificity-determining residues within peptides that are typically between 3 and 11 amino acids in length and can be described by regular expression patterns. Due to the limited number of specificity determinants, novel SLiMs can easily evolve *de novo*, adding new functionality to proteins [2]. Motif binding strongly depends on the context, e.g., functional instances mainly occur inside intrinsically disordered regions (IDRs) that are accessible for interaction [3,4].

Recently, together with publications focusing on the motif sequence, the number of articles dealing with the possible contribution of the disordered flanking regions to motif function is rising. For instance, Stein and Aloy have shown that the contextual contribution to the binding energy of peptide-based interactions is 20% on average. Their results suggest that the context plays a crucial role in maximising binding specificity [5]. Furthermore, coarse-grained simulations have shown that aggregation of small hydrophobic binding motifs can be suppressed by embedding the motifs in disordered regions that are able to sterically stabilise the peptides and hinder the formation of aggregates [6]. In the case of SH2 domain binding motifs Liu *et al*. have found that the contextual specificity is crucial for directing phosphotyrosine signalling [7]. Moreover, Kelil *et al.* have found that although in the case of predicted SH3-binding sites the amino acid sequences are highly conserved compared with their flanking sequences, the latter segments also have a determinant role in positive/negative binding selectivity [8].

Besides some observations related to the probable significance of motif flanking regions in general, there are also examples on how these segments can specifically fine-tune motif binding. One way is to potentially increase affinity via additional binding positions. This is the case during the interaction of the nuclear coactivator binding domain (NCBD) of the CREB binding protein (CBP) and the CBP interaction domain (CID) of the p160 transcriptional co-activator NCOA3, where the binding of the relatively long interaction motif of CID is promoted by short-lived non-specific hydrophobic and/or polar contacts between the flanking regions and NCBD [9]. It has also been shown that the enterohaemorrhagic *Escherichia coli* (EHEC) protein EspFU provides a superior binding affinity compared to cellular ligands of the SH3 domain of the host insulin receptor tyrosine kinase (IRTKS) by utilizing a tryptophan switch in the tripeptide linker between its two PxxP motifs [10]. Another example is the proliferating cell nuclear antigen (PCNA), a cellular hub protein in DNA replication and repair, which is also a potential anti-cancer target. Several proteins interact with PCNA via a SLiM known as the PCNA-interacting protein-box (PIP-box). A systematic study has revealed that the PIP-box affinity can be modulated over four orders of magnitude by additional positively charged residues in the flanking regions [11]. Recently, a new screening method uncovered examples on how the context can influence SLiM’s binding to the EVH1 domain of the cytoskeleton regulator ENAH, that is highly expressed in invasive cancers [12]. In addition, the presence of aromatic residues directly flanking a SLiM in the actin bundling protein Drebrin prevents its interaction with the scaffold protein Homer, which interaction is likely involved in the modulation of synaptic actin cytoskeletons. This example shows that SLiM sequence context can also inhibit motif-based interactions [13]. Furthermore, it has been suggested that the flanking region of radical induced cell death 1 (RCD1)-binding SLiM of DREB2A (Dehydration responsive element binding-protein, a transcription factor known to be involved in plant stress responses) contributes to the stabilisation of the complex possibly through enthalpy-entropy compensation [14].

Many viral proteins use SLiMs in host–pathogen interactions that resemble motifs carried by host proteins. This phenomenon is called linear motif mimicry and helps viruses or other pathogens hijack and deregulate host cellular pathways [15]. These pathogenic SLiMs also often use additional binding sites in their flanking regions to overcome their host counterparts. For instance, the Retinoblastoma protein (pRb)-binding LxCxE and pRb AB groove SLiMs contain charged amino acid stretches in their flanking segments and thereby fine-tune their binding affinity and specificity [16,17]. In addition, phosphorylation of flanking residues can also regulate LxCxE motif binding [18].

In many cases, SLiMs can adopt a well-defined structure before or during partner binding that is also largely influenced by flanking residues. The length and stability of these transient secondary structural elements is crucial for proper complex formation and the mutation of even a single position can have a dramatic effect on the structure and consequently on the function as well. Helix-capping prolines can influence residual helicity and binding affinity by controlling the length of the helix, thus the lifetime of the bound complex. This has been shown in the case of the MDM2-binding motif of the p53 protein, where the proline 27 to alanine (P27A) mutation increases the residual helicity and causes a 10-fold increase in affinity for its ordered binding partner, MDM2. This high-affinity binding to MDM2 is associated with short and weak pulses of p53 activity and reduced p21 expression [19]. However, mutation of helix-flanking prolines to alanines is not always associated with an increase in affinity, as a significant decrease was observed for MLL:KIX, presumably due to the influence of the Pro residue on the interaction of the preceding Leu side chain [20]. The helix initiating and terminating role of helix-capping prolines has also been demonstrated by molecular dynamics (MD) simulations of pre-structured motifs (PreSMos) [21]. Furthermore, it has been suggested that the presence of proline residues adjacent to the RGD motif in dendroaspin and other venom proteins may provide a favourable conformation of the solvent-exposed RGD site for its interaction with integrin receptors [22].

The above findings show that beyond flexibility and accessibility in general, utilizing the disordered context seems to be a perfect way to specifically fine-tune the motif’s binding properties through contributing to its specificity and binding affinity, or influencing its regulation and even its structure. It is still unclear, however, if these segments share any properties in common, which would be indispensable for the emergence of novel functional motifs in a highly variable environment. Such additional information besides the very short and often degenerate regular expressions would make the prediction of these binding sites more accurate as well. In the case of IDRs, Zarin et. al have already shown using an evolutionary approach that most disordered regions contain multiple molecular features that are under selection and that IDRs with similar evolutionary signatures are capable of rescuing function *in vivo*. Their results indicate that sequence-based prediction of IDR functions should be possible based on their physico-chemical properties [23].

Thus, to shed light on the possible role of hitherto unrecognised properties in the function of SLiMs and their surrounding disordered regions, we collected all the known experimentally validated ligand-binding motifs and described them and their flanking regions with different amino acid indices. Our main goal was to find general attributes which need to be preserved within the disordered flanking regions of SLiMs for proper functioning. The results of the present bioinformatics study demonstrate that despite the high sequence diversity within the motif context in general, some restrictions can be found regarding the composition of the flanking regions in several motif classes and in higher motif categories as well.

## Results

To find so far hidden properties and function-determining factors in the known domain-binding motifs and their flanking regions we started with developing a new approach that could help us further understand their features and characteristics. Motifs are usually embedded in IDRs and it has already been shown that around 20 residues can be considered as unstructured on both sides [3]. Based on earlier findings in the field of SLiMs (see Introduction) we propose that the function-determining factors within their flanking regions are usually not position-specific and cannot be captured by examining the conservation of the amino acid sequence itself. In addition, the determination of motif boundaries can also be challenging in many cases since they can be extremely difficult to define even with experimental methods. Therefore, we aimed to investigate different parts of the flanking regions within 20 residues from the motif boundaries instead of specific positions. We also suggest, based on previous findings in the field of IDRs that the composition and the physico-chemical properties of these protein segments must be the key to find additional information on their function. Thus, we have decided to examine the motifs and their flanking regions based on functionally important amino acid properties / indices. To reduce complexity and to ensure that the sequences could be comparable, we have used simplified numbers representing the original indices of each amino acid. The mean values of different flanking segments have also been calculated to capture their basic characteristics.

### Characterisation of the flanking regions by simplified amino acid properties

We performed a property-based bioinformatics analysis on the flanking regions of domain-binding motifs, where we focused on the attribute diversity and restrictions in 6 different parts of these segments. To this end, we retrieved all the known experimentally validated general ligand-binding sites from the ELM (true positive LIG, DEG and DOC motif instances) and LMPID databases. Both databases contain manually curated motifs from the literature. Although ELM is the most widely used resource of SLiMs and is also more regularly updated compared to LMPID, with careful reconciliation of motif classes and boundaries the latter could perfectly complement our dataset. For the property-based analysis we decided to characterise the disordered flanking segments of motifs with 8 biologically relevant amino acid indices that reliably represent their physical, chemical or structural characteristics and are not highly correlated with each other: Kyte-Doolittle hydropathy, volume, isoelectric point, charge, negative charge, positive charge, proline content and serine/threonine content. Hydropathy, net charge and volume are attributes that are known to have the ability to distinguish between order and disorder [24,25] tendencies. Since flanking regions have considerably less charged residues than IDRs [3], we used charge content instead of net charge in this work for the comparison of different parts of the flanking regions. In addition, proline-rich segments, highly negatively and positively charged sequences [26], as well as the number of phosphorylation sites can also be determining for the function within IDRs [27]. To create a robust and comparable dataset, the residues of the motifs and their 20 residue-long flanking segments were substituted with numbers (1 to 5) representing the linearly rescaled AAindex values of each amino acid for a given feature (see Methods, S1 Appendix and S2 Appendix). Since this work addresses characteristic physico-chemical features of SLiMs and their flanking regions, we divided the flanking regions into smaller, 5 and 10 residues long segments and performed calculations and comparisons on these regions separately. Through this approach, the simplified properties of each part of the flanking regions could be determined and further examined (Fig 1).

**Fig 1.**
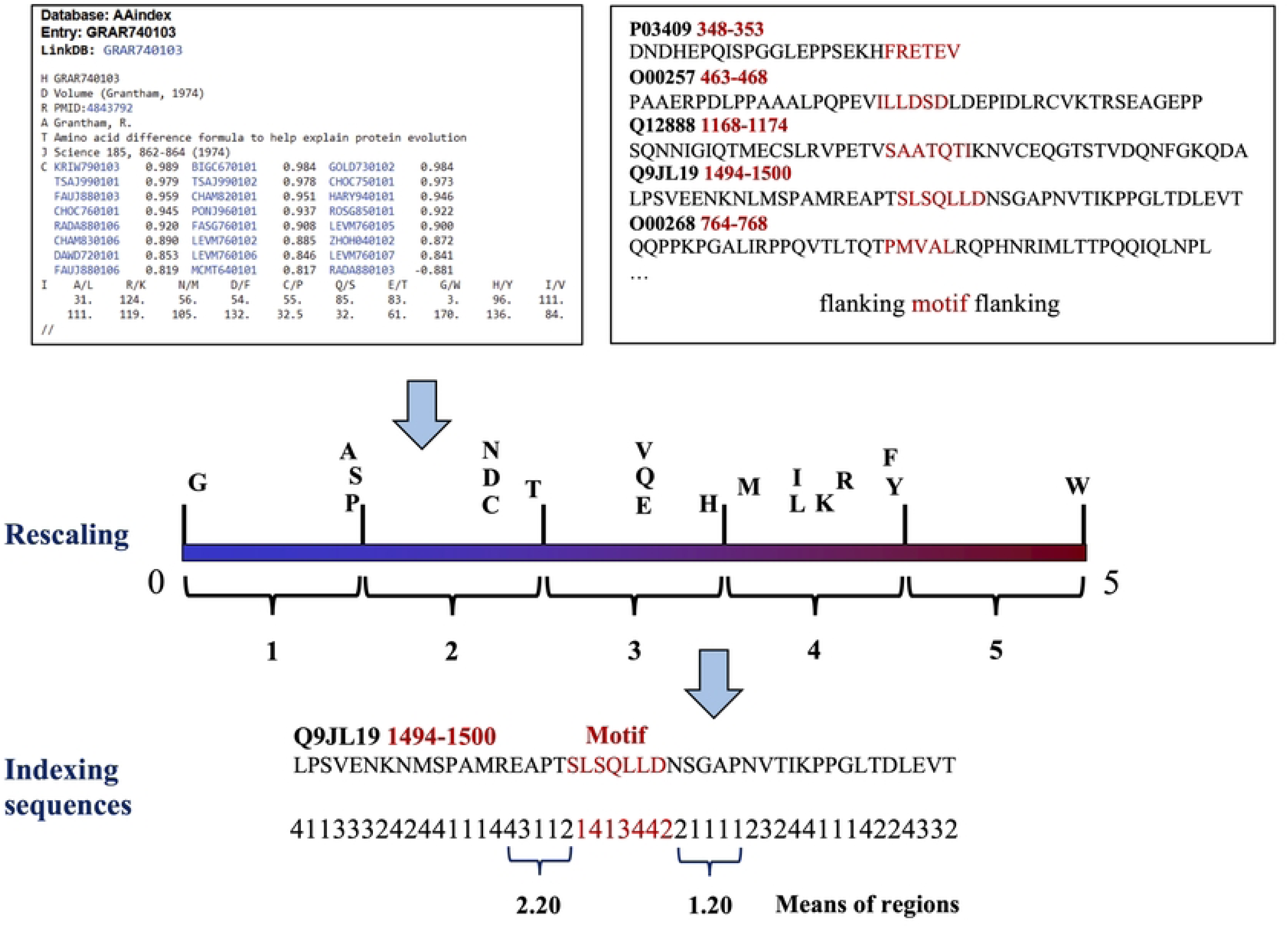
Schematic representation of the indexing process using amino acid size as an example feature.

### The flanking regions show higher property diversity than the motifs

In the ELM database, SLiMs are categorised into motif classes, where instances share function or partner domain and are described with the same regular expression [28]. The motif core – which is sufficient for partner binding – is defined by fixed and degenerate positions (where none, or only limited types of substitutions are tolerated) and additional sequential restrictions (like prohibited residues in certain positions) that are specific to each class and provide relevant information for motif searching methods. Here, the ranges (largest differences) of the property mean values of the motifs and their 5 AA-long N- and C-terminal flanking segments (N5 and C5, respectively) were calculated for 42 motif classes (classes that contain at least 10 motif instances in our dataset) (S1 Table). From these data, 6 attributes of 10 randomly selected representative motif classes are shown in Fig 2A. We found that the flanking regions tend to show higher diversity for all investigated features, compared to the motifs. To confirm this finding statistically, we performed paired t-tests using the ranges of the 42 large classes. We found that both the N5 and C5 regions show significantly higher diversity than the motifs (the p-values were <1.23E-07 for all the 8 properties) (S2 Table). Moreover, in the case of the 10 randomly selected classes, the difference between the maximum and minimum mean values of the motifs is at most 2.0, while in the 5 residue-long flanking sequences goes up to 4.0. However, we also observed in a few cases that the motifs also show relatively high variability (e.g., the proline content of the LIG_SH3_3 and DOC_WW_Pin1_4 classes, the Ser/Thr content of DOC_WW_Pin1_4 class, or the charge content of LIG_SH2_CRK and LIG_SUMO_SIM_anti_2 classes (S1 Table).

**Fig 2.**
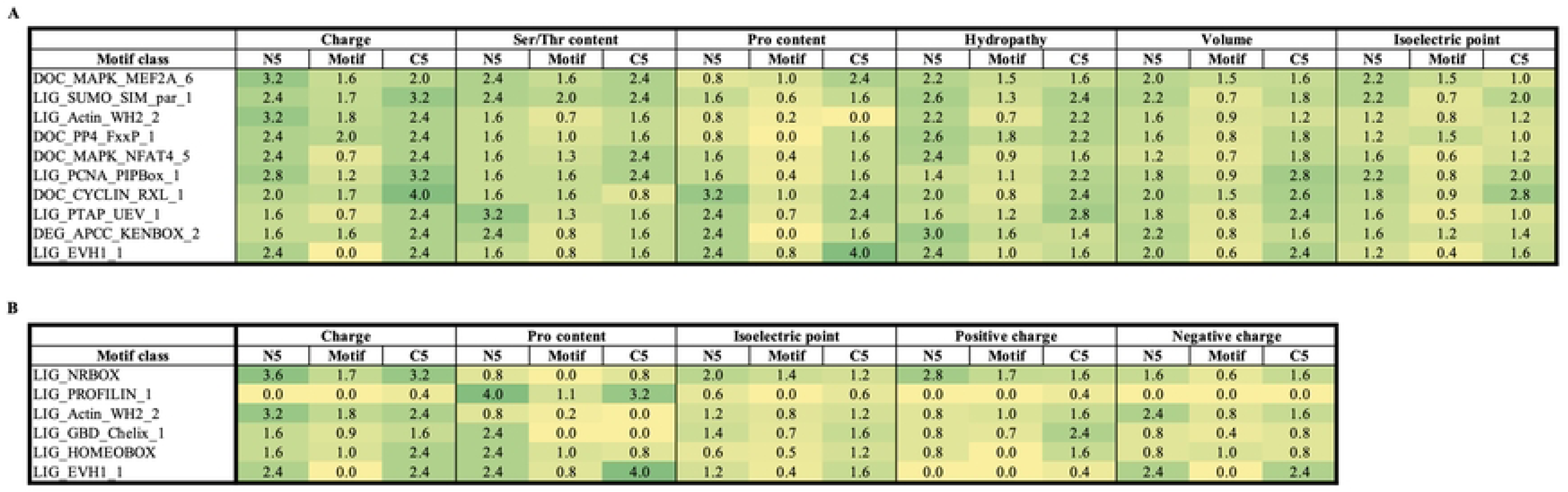
Variability of properties within the motifs and their flanking regions. N5: first 5 residues upstream of the motif, C5: first 5 residues downstream of the motif. A: Largest ranges of mean values within 10 randomly selected motif classes are shown. B: Largest ranges of mean values of 6 motif classes where small (≤ 0.8) variety can be found within both flanking regions. Darker green colours represent higher variations, yellow colours represent smaller variations.

In the case of proline, serine/threonine and charge indices, if the contents of two sequences differ by only one amino acid (e.g., 0 or 1 proline is present) within a 5 residue-long region, the difference between the mean values will be 0.8. Thus, a maximum value of 0.8 is considered a small range, while a value of > 0.8 is considered a high range (maximum – minimum mean value of a region) within a motif class. Analysing the ranges using this threshold we observed that in some cases flanking segments also show low variety, especially in their proline content or negative/positive charge contents (S1 Table). These results indicate that in some motif classes flanking regions close to the motif core may also have features that need to be under evolutionary constraint. In this first analysis we filtered the motif flanking regions to be highly disordered (see Methods) to ensure that potentially structured flanking regions (that may influence the binding properties and transient structure of the motif) are not included. In the next section we extend the scope for all motifs and flanking regions to validate if these findings are generally applicable to certain ELM classes.

### Several motif classes show low property variety close to the motif core

We repeated the previous analysis with all the available motif instances of the ELM database (version of February 2023, without disorder filtering) in the case of those motif classes, where we earlier found a property with low variety (a maximum of 0.8 range) within at least one of their flanking regions. We found 19 elm classes where at least one of the investigated amino acid properties were conserved within the 5-residue long flanking regions (Table 1). Furthermore, in 6 out of the 19 classes we observed small variations that were symmetrical (the ranges are small on both sides of the motifs) (Fig 2B). Overall, the LIG_PROFILIN motif class shows the highest conservation of charge properties in both flanking regions due to the complete lack of charged residues adjacent to its motif instances. These proline-rich SLiMs bind to a hydrophobic groove of profilin, a key regulator in actin polymerization, and based on our results, charged side chains are not tolerated for this interaction. LIG_PROFILIN motifs are often involved in multiple binding, while their flanking regions are also enriched in Gly and Pro residues, thus, extreme flexibility may also be crucial for their functions. LIG_Actin_WH2_2 motifs also contribute to actin filament assembly through binding to G-actin. They all consist of an N-terminal short helical, and a C-terminal disordered region. We found that the proline content of the N5 and C5 flaking regions of WH2 SLiMs is generally low (0 or 1 Pro occurs in these sequences). Similarly, the nuclear receptor (NR)-binding LIG_NRBOX motifs adopt helical structures upon binding and based on our findings, a maximum of 1 proline residue is tolerated within their proximity. The 5 residue-long flanking regions in the GTPase-binding domain (GBD) ligand LIG_GBD_Chelix_1 motif class also seem to be under constraint. Both N5 and C5 contain at most 1 negatively charged residue, while the mean hydropathy of N5 and the volume, Pro content and Ser/Thr content of C5 all show very low diversity among the instances of the motif class. In the class LIG_KLC1_WD_1 – which binds to the TPR domain of kinesin light chain 1 and is involved in cellular cargo transport –, a maximum of 1 positively charged residue can be found in the N-terminal flanking regions of the instances. This is in accord with the fact that the surface of the binding groove of the TPR domain is positively charged which can ideally stabilise the acidic binding motif through electrostatic interactions [29].

**Table 1.** Conserved properties in the flanking regions. ELM classes, where certain AA properties of the 5 residue-long flanking regions are preserved are shown (the difference between the maximum and minimum mean values is ≤ 0.8).

|  | **N5** | **C5** |
| --- | --- | --- |
| **Charge** | LIG_PROFILIN_1 | LIG_PROFILIN_1 |
| **Ser/Thr content** |  | LIG_GBD_Chelix_1 |
| **Pro content** | LIG_NRBOX<br>LIG_Actin_WH2_2<br>DOC_PP4_FxxP_1 | LIG_NRBOX<br>LIG_Actin_WH2_2<br>LIG_GBD_Chelix_1 |
| **Hydropathy** | LIG_GBD_Chelix_1 |  |
| **Volume** |  | LIG_GBD_Chelix_1 |
| **Isoelectric point** | LIG_PROFILIN_1<br>LIG_HOMEBOX | LIG_PROFILIN_1 |
| **Positive charge** | LIG_EVH1_1<br>DOC_PP4_FxxP_1<br>LIG_PROFILIN_1<br>LIG_HOMEBOX<br>LIG_KLC1_WD_1<br>LIG_CtBP_PxDLS_1<br>LIG_WRPW_1<br>LIG_SUMO_SIM_anti_2 | DOC_SPAK_OSR1_1<br>LIG_EVH1_1<br>LIG_PROFILIN_1<br>DOC_MAPK_NFAT4_5 |
| **Negative charge** | LIG_PROFILIN_1<br>LIG_GBD_Chelix_1<br>LIG_HOMEBOX<br>DOC_MAPK_JIP1_4<br>LIG_SH3_1 | LIG_WW_1<br>LIG_CSL_BTD_1<br>LIG_PROFILIN_1<br>LIG_GBD_Chelix_1<br>LIG_HOMEBOX<br>LIG_CtBP_PxDLS_1<br>LIG_PCNA_PIPBox_1<br>LIG_PAM2_1 |

### The percentage distribution of properties is similar in different parts of the flanking regions

To find out if there are any differences between the frequencies of properties in different parts of the flanking regions, we calculated the percentage distribution of 5 simplified mean values in 6 parts of our sequences (Fig 3). This analysis was performed on 20×120 random samples from our nonredundant and nonbiased collection (1192 motifs, see Methods). Here, besides the simplified low (the mean value of the region is >1 and ≤ 2), medium (the mean value is >2 and ≤ 3), high (the mean value is >3 and ≤ 4), and very high (the mean value is >4) categories, we also included ‘none’ mean value indicating that such an attribute is not present within a protein region (the mean value is 1). We found that different parts of the context usually show very similar properties in the N- and C-terminal flanking regions, and in most cases, in the first and second 10 residue-long flanking regions (regions 1-10 and 11-20, respectively) as well. Although no remarkable differences could be detected comparing these regions, some characteristic preferences could be established for the flanking segments. For example, regarding the Kyte-Doolittle hydropathy and isoelectric point, all the 6 investigated regions tend to show intermediate (medium) mean values, sequences with low and high mean values also occur, but the extremities are not represented (Figs 3A and B). For the residue volume index, medium and low mean values occur most frequently, but among the C1-5 regions some show ‘none’ values (where the mean value is 1, since only Ala, Gly, Pro or Ser residues occur within the sequence). Not surprisingly, there are no regions with very high mean side chain volume values amongst our disordered flanking sequences as high-volume residues typically have large hydrophobic side chains (Fig 3C).

**Fig 3.**
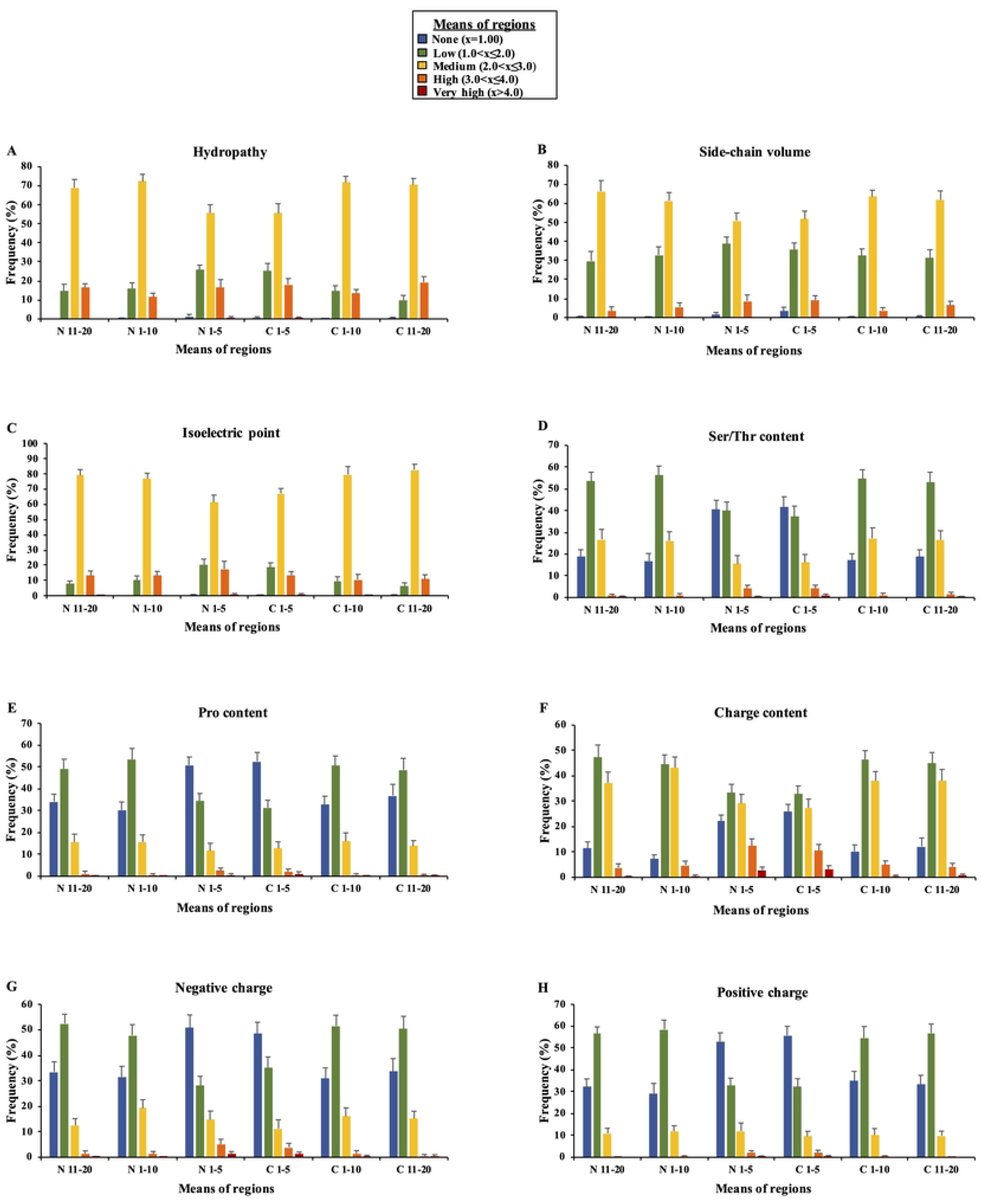
The frequencies of property means in different parts of the flanking regions. (1192 motifs, random 20×120 sequences, 80% maximum similarity). N: N-terminal flanking, C: C-terminal flanking.

Focusing on the Ser/Thr content, in N1-5 and C1-5 regions, ‘none’ and low mean values are the most common. High and very high Ser/Thr content is rare in all regions, while there are no sequences with very high values within the first 10 residue-long flanking segments (N1-10 and C1-10) (Fig 3D).

Proline content is also very similar in the N-and C-terminal flanking regions. More than 50% of the sequences do not contain any proline residues within 5 amino acids, while high and very high Pro content is extremely rare. In N1-10 very high means of Pro content are completely missing (Fig 3E).

The highest variability can be found in the case of charge content, where low and medium mean values are relatively common within N1-5 and C1-5, but highly charged sequences also occur. In the 10 residue-long flanking regions most sequences have low or medium mean values, but highly charged and uncharged sequences are also present in our dataset. Interestingly, the mostly charged region is N1-10, where we can see some slight differences (’none’ mean charge is less frequent, while medium is more frequent) comparing with the other parts of the flanking regions (Fig 3F). Within N1-5 and C1-5, most sequences do not have positively charged residues, while it is rare that motifs are embedded in a highly positively charged protein region. Interestingly, very high mean values cannot be found in the 10 amino acid-long segments of the flanking regions at all, indicating that too many positively charged residues close to each other are not favourable in the motifs’ proximity (Fig 3G). Negative charge content shows very similar tendencies to positive charge content, except that medium, high and very high mean values all show higher frequencies in the former case (Fig 3H).

### Specific compositional biases in the different ELM category motif flanking regions

Since the 2014 release of ELM, docking (DOC) and degradation (degron, DEG) motifs have been classified separately from the classical ligand-binding (LIG) sites [2]. Although all three types are ligands of globular domains, docking sites typically interact with kinases and phosphatases separate from the active site [30], while DEG motifs target the protein for degradation, and several mutations in their flanking segments have been linked with decreased degradation rates and severe diseases [31]. To see if there are any differences in the distribution of properties between the 3 separated subcategories, we repeated the previous analysis with smaller datasets including only LIG, DEG or DOC motifs (S1 Fig). Only slight differences can be detected in the distribution of flanking properties comparing the 3 subcategories. The most remarkable finding is that in at least one of the 3 investigated categories some ranges of mean property values that could theoretically occur within IDRs are not found at all. For instance, among the flanking regions N1-10 and C1-10 of DEG type motif classes there are no sequences with very high charge content, while the same flanking regions of certain LIG and DOC type motifs are highly charged. The 10 residue-long flanking regions of degrons also seem to lack sequences with very high proline content (this is also true for DOC N1-10, N11-20, and C1-10), while there are several LIG motifs in our dataset that are surrounded by many proline residues.

To see what properties are completely missing in the sequence context of all the currently known ligand-binding SLiMs, we collected all motifs and their flanking segments from the latest version of the ELM database (last modified on: March 14, 2023) and repeated the analysis without redundancy and disorder filtering. The occurrence of sequences with extreme composition (‘none’, high and very high mean values) in the LIG/DEG/DOC datasets is summarized in Fig 4. Here, the most interesting observation regarding all ligand-binding motifs (all the 3 subcategories) is that very high positive charge mean values are completely missing in the 10 residue-long flanking segments, while very high negative charge can occur in some regions. While there are no sequences with very high negative charge in the N-terminal flanking segments of the DEG and DOC motifs, SLiMs of the LIG subcategory can be highly negatively charged. Apparently, unlike DEG and DOC motifs, LIG sequences can be embedded in a more structured or an extremely flexible environment as well, since very high mean hydropathy values and ‘none’ mean volume values (where only tiny residues are present) are both tolerated by functional LIG SLiMs. Furthermore, LIG, DOC and DEG SLiMs all have flanking segments where very high Ser/Thr content does not occur. Degrons seem to have the most restrictions regarding the composition of their flanking regions. Unlike LIG motifs and docking sites, they are not located in regions where the 10 residue-long flanking segments have very high proline content, or the first 10 residues are extremely charged.

**Fig 4.**
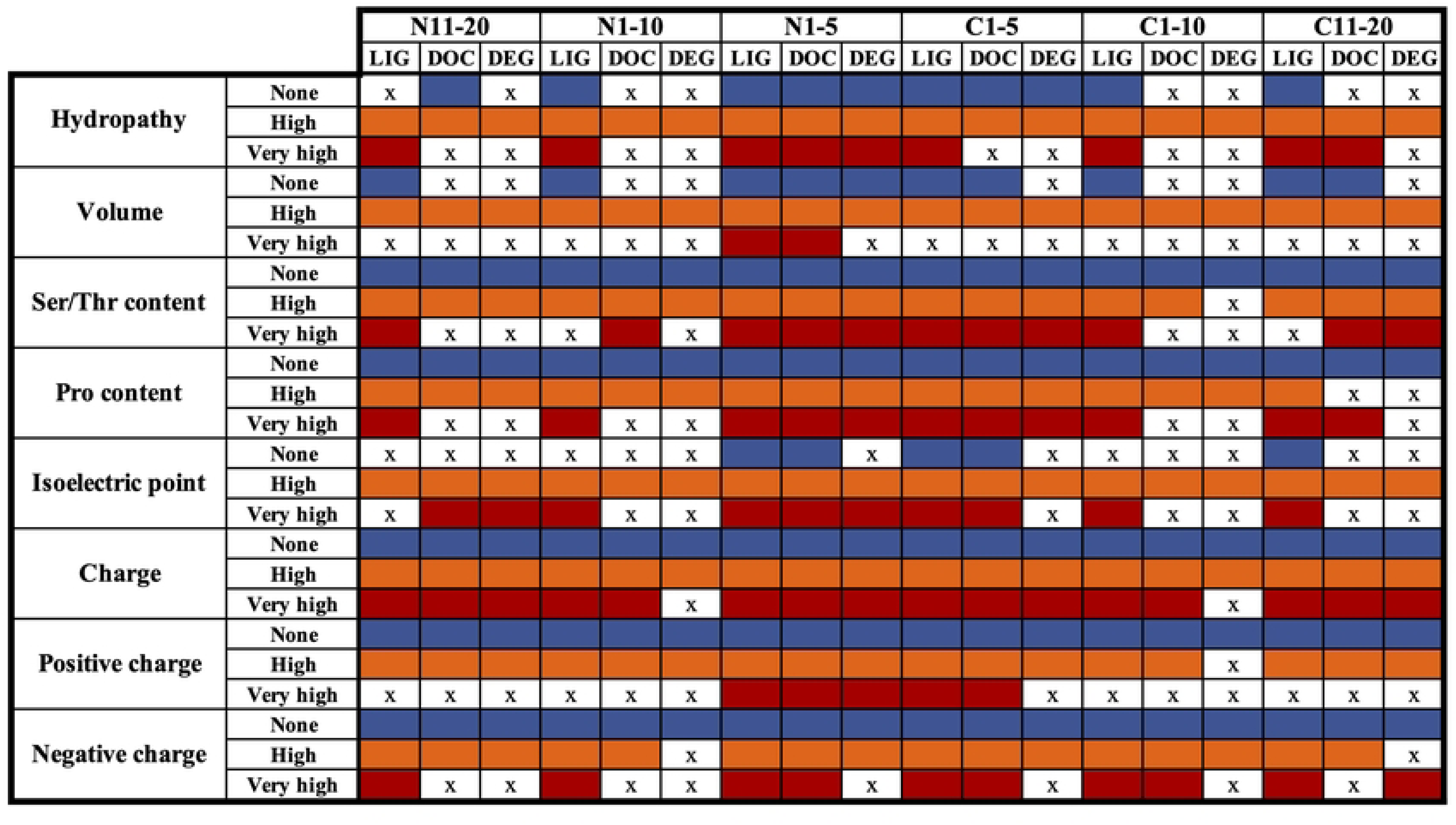
Occurrence of ‘none’, ‘high’ and ‘very high’ mean values in the LIG/DOC/DEG flanking regions. The coloured cells indicate that sequences with such mean value are present in the dataset, while ‘x’ denotes that there are no sequences with such property in the given category.

### Overlapping and adjacent motifs have smaller means of volume in general

It has been already shown that SLiMs are abundant in hydrophobic and charged residues (which is particularly important for partner recognition), while are also enriched in floppy and rigid amino acids, which distinguishes them from both globular and disordered protein regions. Flanking regions, however, are depleted in hydrophobic residues and contain less charged amino acids than generic IDRs [3]. This raises the question whether those motifs that are situated in another motif’s proximity – thus, theoretically need to function as a ligand-binding site and a flexible flanking region of another motif at the same time – show any differences compared to those SLiMs which are ‘alone’ in an unstructured region. To test this, we collected those motifs that are located close to another known, experimentally validated ligand-binding motif (overlapping/adjacent motifs overlap with another LIG/DOC/DEG motif but have at least 4 residues outside the other motif’s core or start within another motif’s 8 residue-long flanking region). As a control dataset we used the remaining of our motif collection, after removing overlapping motifs and close proximity motif incidences (where other known LIG/DEG/DOC motifs can be found within 10 residues from the motif’s boundaries). While we cannot exclude the presence of yet unknown motifs in the flanking regions of these motifs, we believe it is much less enriched and can be used in the comparison. The most remarkable differences could be detected in the case of volume index, where we found that overlapping/adjacent motifs have smaller means of side chain volume in general, compared to the control dataset (Fig 5A). They also use more Ser/Thr and Pro residues, while have less hydrophobic side chains (Figs 5B, C and D). These latter findings also explain the difference in the means of volume index between the two datasets. It is also noteworthy, that in the case of charge content, isoelectric point, and positive/negative charge content separately, we could not detect remarkable differences between the two datasets. Overlapping and adjacent SLiMs can derive from the same, or even from different motif classes, and there are many examples in the literature on how they can influence each other (e.g., co-operatively or even competitively) [32]. Based on these data it is plausible that the usage of more small/tiny residues in the case of overlapping/adjacent SLiMs is necessary to avoid steric interference between the motifs as well as to retain local flexibility.

**Fig 5.**
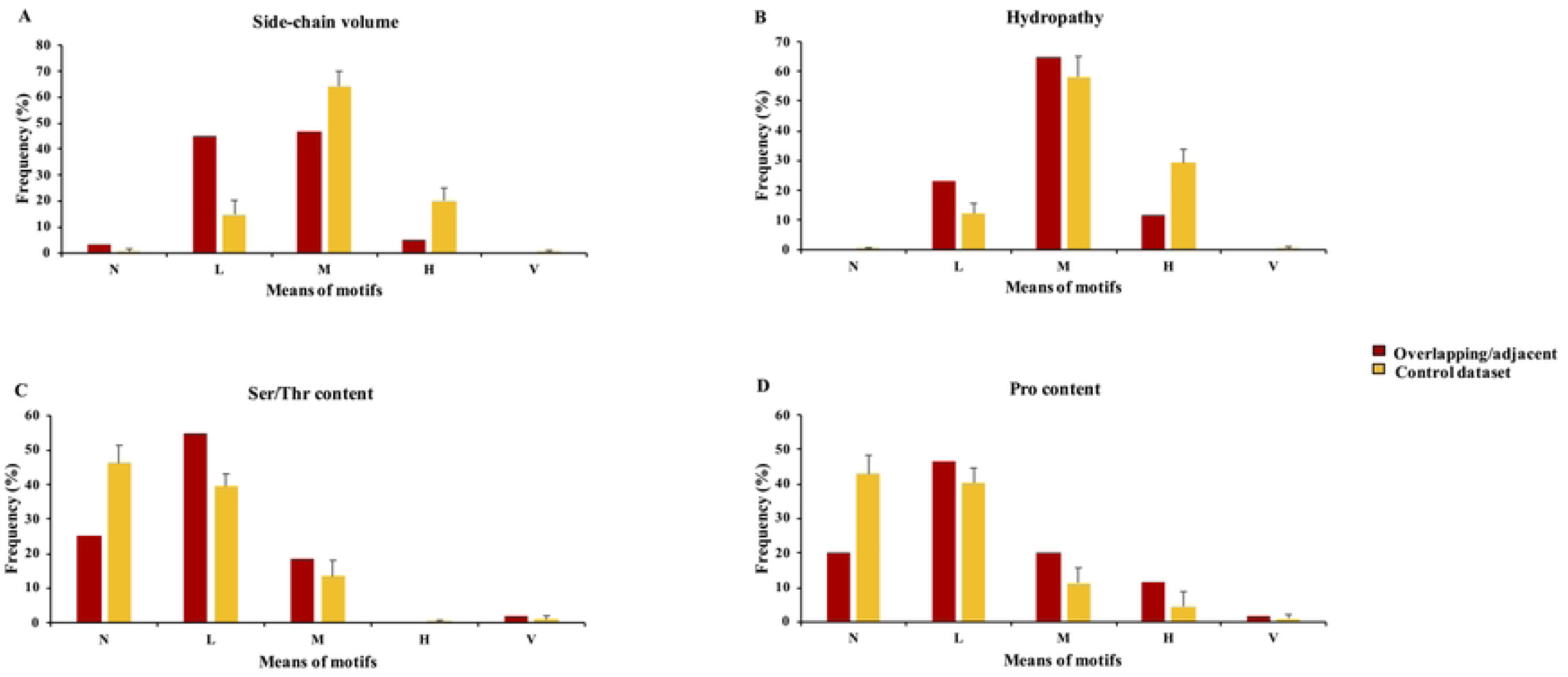
The frequencies of property mean values of overlapping/adjacent motifs and the control dataset. N: None, L: Low, M: Medium, H: High and V: Very high.

## Discussion

Earlier, the work of Chica *et al.* already indicated that the prediction of linear motifs can be complemented with contextual information [33]. Conversely, Davey *et al.* have found that functional SLiMs show higher levels of conservation than their context, and that flanking regions of targeting and ligand binding sites most closely resemble the IDRs based on amino acid attributes of the sequences [34]. In the present study we focused on the potential contribution of the flanking regions to motif function in general. We investigated the diversity, property distribution, and preserved features in the flanking regions of the 3 ligand-binding categories of SLiMs (LIG, DEG and DOC) and 42 large motif classes of the ELM database. We found that in many cases, additional information can be captured on the feature level in the flanking segments that might be under evolutionary constraint and contribute to proper motif structure and function.

By comparing the ranges of index mean values of the motifs and their 5 residue-long flanking regions within larger ELM classes we found that the flanking segments show higher diversity than the motifs in general (even though motifs usually contain only a few fixed positions and considered as degenerate segments). Such a high level of diversity can give potential to specifically fine-tune each individual motif’s binding properties, as exemplified by the cases described in the ‘Introduction’ section. Yet, we found 19 classes where at least 1 index shows very low variety and thus seems to be evolutionarily preserved. In class level categories (ligand-binding sites in general, and LIG/DEG/DOC motifs separately), we also found that while the property content is well-distributed in all parts of the flanking regions, some types of IDRs are not represented. For example, very high positive charge content is not present in the 10 residue-long flanking regions of the examined ligand-binding motifs. The reason of this observation is unclear at present, but we speculate that while a limited number of additional positively charged amino acids could specifically fine-tune motif binding, many of them would result in undesired interactions in the cell (e.g., DNA or RNA binding). Furthermore, LIG, DOC and DEG SLiMs all have flanking segments where very high Ser/Thr contents never occur. A recent review demonstrates how phosphorylation events on these side chains can modulate the motif’s binding affinity [35], thus it is reasonable to think that there must be some constraint on the number of Ser/Thr residues in these regions.

Out of the 3 categories, degrons seem to have the most restrictions regarding the composition of their surrounding regions. DEG motifs do not have very highly charged N1-10 and C1-10 sequences (or very high negative charges in 3 regions), and they do not use very high Ser/Thr content in 3 out of 4 10 residue-long flanking segments, either. It has recently been shown that additional Ser, Thr, or Tyr residues surrounding DEG motifs can regulate or even rescue degron function [31]. However, regions that are abundant in serines can be highly modified (e.g., phosphorylated or glycosylated), and extremely high serine content has been linked to various specific regulatory processes [36]. Thus, it seems plausible that the motif context does not prefer poly-Ser/Thr sequences in order to avoid unfavourable interactions / modifications. Similar observation was made with the Pro index of the 10 residue-long flanking regions of DEG (and 3 out of 4 flanking regions of DOC) motifs, where no sequences with very high Pro content could be detected. Pro-rich disordered segments often promote polyproline type II (PPII) helix conformations [37], that would likely provide an environment that is too rigid for the proper functioning of these types of SLiMs, while they could also be involved in additional hydrophobic interactions.

Our findings indicate that positive charge content, and in some regions, negative charge, Pro and Ser/Thr content all need to be under evolutionary constraint in the context of SLiMs. Kastano *et al.* have recently shown that Ser, Pro, Glu and Lys are the most abundant residues within compositionally biased regions (CBRs) of the human proteome. However, if we focus on CBRs located in IDRs involved in hub interaction networks, Ser-, Pro- and Glu-rich regions are the most common, while K-rich regions barely occur [38]. The enrichment of Ser and Pro residues in CBRs can be explained by the presence of tandem SLiMs generated for cooperative regulation, that is in line with our findings regarding overlapping/adjacent motifs. It is particularly interesting, though, that while Ser-, Pro- and Glu-rich segments are abundant in CBRs, motif flanking regions seem to avoid the usage of these types of disordered regions, presumably due to their abovementioned involvement in regulation and additional protein– protein interactions.

SLiMs are important and highly abundant [39] protein functional modules. However, certain features of the motifs cannot be efficiently captured by the regular expressions, especially those that cannot be rendered to specific positions. Although we cannot rule out the possibility that some of the examined classes and higher categories would allow wider variety of the features we found to be preserved, the above findings all indicate that the examination of disordered sequences based on amino acid properties is a valid approach to reveal important features in the flanking regions of SLiMs that are otherwise barely recognisable. Moreover, since the computational identification of motifs is hindered by their compactness and the relatively low numbers of well-defined positions, preserved features in the flanking regions can also add further information to motif searching and prediction methods. In addition, our finding that overlapping/adjacent SLiMs seem to use smaller side chains could also be taken into consideration during the identification of new motifs located in other motifs’ proximity. Our approach provided further information on the characteristics of the disordered flanking regions of domain-binding motifs that can help us understand motif evolution better, while it can also give potential to expand motif searching and prediction methods.

## Methods

### Dataset generation

All the known experimentally validated, true positive motif instances were downloaded from the Eukaryotic Linear Motif (ELM) database [40] (http://elm.eu.org, version of 15.10.2020). In this work, we aimed to examine those SLiMs that are ligands of globular protein domains, thus only the LIG/DEG/DOC motifs were selected for further investigation. SLiMs from the LMPID database [41] (http://bicresources.jcbose.ac.in/ssaha4/lmpid/) were also downloaded and added to the dataset. Next, we performed a preliminary redundancy filtering, where identical motif regions within the same proteins (identical UniProt ID and motif boundaries) were filtered out. Protein sequences were downloaded from the UniProt database (https://www.uniprot.org/) and motifs with their 20 residue-long flanking regions were collected. Disorder filtering of the motifs and their surrounding segments were performed using the IUPred2A long algorithm [42] (https://iupred2a.elte.hu/). Since our main purpose was to investigate the features of SLiMs located in highly flexible and variable protein regions, we tried to filter out sequences where at least one of the flanking segments would fall into a more structured region. Based on our comparison of different filtering settings, many sequences where at least 50% of the residues have IUPred scores ≥ 0.4 failed to satisfy this criterion. Thus, sequences where at least 50% of the residues have scores ≥ 0.5 were considered disordered and kept in the dataset. Also, because our dataset still contained some motifs based on only prediction methods, those instances were retained which were supported by at least one experimental evidence. Finally, we applied a second redundancy filtering for significantly overlapping motifs (the motif boundaries differed in 2 residues at most). We also compared the motif classes of the ELM and LMPID databases and ignored those from the latter, which did not ‘fit in’ one of the known ELM classes (or the motif boundaries were not identical to the boundaries of the equivalent ELM class). The only exception was the LIG_Rb_LxCxE_1 class, where the true length of the motif is arguable based on the large difference between the boundaries in the two databases we used. Thus, in this case we changed the lengths of the motif instances to 8 residues in all cases. The final dataset contained 1297 ligand-binding motifs and their 20 residue-long flanking regions.

Our dataset contained 3 large motif classes (LIG_14-3-3_CanoR_1, DOC_WW_PIN_1_4 and LIG_EH_1) with 60-80 instances in each. In order to avoid any bias, for the analysis of the percentage distribution of the property mean values in the flanking regions and within nonoverlapping motifs (see below), we randomly decreased the number of motifs from these classes to 30, as all the other classes contained approximately 10-30 instances.

### Selection of amino acid properties

Amino acid indices were downloaded from the Amino acid index (AAindex) database [43] (https://www.genome.jp/aaindex/) and 3 biologically relevant properties were chosen for our analysis: Kyte-Doolittle hydropathy, volume and isoelectric point (AAindex accession number KYTJ820101, GRAR740103, and ZIMJ680104, respectively). We also added 5 further attributes representing different amino acid contents typically enriched in disordered protein regions: charge, negative charge, positive charge, proline content and serine/threonine content.

### Rescaling, indexing sequences

Since not only the amino acid values but also the scale of the indices in the AAindex database are very diverse due to the different methods the authors used, we needed to reduce this complexity to simplify and also rescale our data. For this purpose, first we divided the scales to 5 equal parts, and placed the amino acids on them according to their original values of properties (Fig 1, S1 Appendix). Then, we substituted the side chains with numbers 1 to 5 based on their location along our simplified scale. Here, amino acids with the lowest values were replaced by number 1, while those having the highest values were substituted with number 5. In the case of charge, proline and serine/threonine content indices we only used numbers 1 and 5 as absent or present respectively (except for positive charge, where His received number 3).

Following this, we substituted all the amino acids with numbers 1 to 5 along the collected motif sequences and their flanking regions. This indexing step was performed with every property we selected for this work (S2 Appendix). Mean values of the motifs and their 5, 10 and 20 amino acid-long flanking regions were calculated for the investigation of the characteristics of the substituted sequences. Sequences that did not contain all residues of an examined flanking segment were omitted from the calculation of mean values.

### Paired t-tests

Shapiro-Wilk normality tests and paired t-tests were performed for both the N5-motif and C5-motif range value pairs of the 42 large ELM motif classes using the R program (http://www. r-project.org/).

### Distribution of property mean values in different parts of the flanking regions

To analyse property characteristics in different parts of the flanking regions, first we simplified the calculated mean values by substituting them with one of 5 categories (‘none’: only number 1s occur in the region, low: the mean value is >1 and ≤ 2, medium: the mean value is >2 and ≤ 3, high: the mean value is >3 and ≤ 4, and very high: the mean value is >4. Next, we generated 20×120 random datasets from our nonbiased dataset (1192 motifs) and performed a redundancy filtering using the online Expasy decrease redundancy program (https://web.expasy.org/decrease_redundancy/). Here, sequences (motifs with their 5-5 residue-long flanking regions) with a maximum of 80% similarity were retained in the datasets. The percentage distribution of mean values was calculated using Perl program language.

We repeated this analysis for the LIG/DEG/DOC motifs separately, where the randomized LIG datasets (15×120 out of 854), DOC datasets (6×120 out of 246) and the DEG dataset (88) were also subjected to an 80% redundancy filtering and the percentage distribution of simplified mean values were calculated for 6 different parts of the flanking regions.

### Investigation of all LIG/DEG/DOC motifs from the ELM database

We downloaded the latest version of the motif instances dataset from the ELM database (last modified on: March 14, 2023) and collected the ‘true positive’ LIG/DEG/DOC motifs and their 20 residue-long flanking regions in separate datasets. We repeated the indexing process with these nonfiltered datasets and calculated the simplified mean values for 6 different parts of the flanking regions (as seen above).

### ‘Overlapping/adjacent’ and control datasets

We collected all motifs from our dataset which contain another ligand-binding motif in their proximity and named them ‘overlapping/adjacent’ motifs (an ‘overlapping/adjacent’ motif overlaps with another LIG/DOC/DEG motif but has at least 4 residues outside the other motif’s core or starts within another motif’s 8 residue-long flanking region). Following a redundancy filtering (80% maximum similarity of the motif sequences and only one motif has been retained from the same class with the same UniProt ID), we created a dataset of 60 nonredundant motif instances. Motifs of the control dataset do not have another LIG/DEG/DOC motif within their 10 residue-long flanking regions. Property means and their ratios of the motifs were calculated (as described above) for the ‘overlapping/adjacent’ and control (10×60 random sequences with a maximum of 80% similarity) datasets.

## Author contributions

Conceptualisation: VA, LK and AT; methodology: VA and LK; research: VA and AH; software: VA and AH; data analysis: VA; resources: AT; data curation: VA; writing - original manuscript: VA; writing – review and editing: LK and AT; supervision: LK and AT; funding acquisition: AT.

## Supporting information

S1 Appendix: Selected indices and the simplified (1 to 5) values of each amino acid.

S2 Appendix: Final dataset of the motifs and their flanking regions containing the original and the indexed sequences.

S1 Table: The ranges (largest differences) of the property mean values of the motifs and their 5 AA-long N- and C-terminal flanking segments in 42 motif classes.

S2 Table: p-values calculated from the paired t-tests showing the significant differences in property diversity between the N5 – motif and motif – C5 regions of 42 motif classes.

S1 Fig: The frequencies of property mean values in different parts of the flanking regions (LIG, DEG and DOC motifs separately). N: N-terminal flanking, C: C-terminal flanking.

